# Genome-wide Association Studies Reveal Similar Genetic Architecture with Shared and Unique QTL for Bacterial Cold Water Disease Resistance in Two Rainbow Trout Breeding Populations

**DOI:** 10.1101/163964

**Authors:** Roger L. Vallejo, Guangtu Gao, Sixin Liu, Breno O. Fragomeni, Alvaro G. Hernandez, James E. Parsons, Kyle E. Martin, Jason P. Evenhuis, Timothy J. Welch, Timothy D. Leeds, Gregory D. Wiens, Yniv Palti

**Affiliations:** National Center for Cool and Cold Water Aquaculture, Agricultural Research Service, United States Department of Agriculture, Kearneysville, WV 25430; Animal and Dairy Science Department, University of Georgia, Athens, GA 30602; High-Throughput Sequencing and Genotyping Unit, Roy J. Carver Biotechnology Center, University of Illinois at Urbana-Champaign, Urbana, IL 61801; Troutlodge, Inc., P.O. Box 1290, Sumner, WA 98390

**Author notes:** **Corresponding author:** Yniv Palti, National Center for Cool and Cold Water Aquaculture, Agricultural Research Service, U.S. Department of Agriculture, 11861 Leetown Rd., Kearneysville, WV 25430, USA.

**Keywords:** Aquaculture, Bacterial cold water disease, Genome-wide association study, Quantitative trait loci, Rainbow trout

## Abstract

Bacterial cold water disease (BCWD) causes significant mortality and economic losses in salmonid aquaculture. In previous studies, we identified moderate-large effect QTL for BCWD resistance in rainbow trout (*Oncorhynchus mykiss*). However, the recent availability of a 57K SNP array and a genome physical map have enabled us to conduct genome-wide association studies (GWAS) that overcome several experimental limitations from our previous work. In the current study, we conducted GWAS for BCWD resistance in two rainbow trout breeding populations using two genotyping platforms, the 57K Affymetrix SNP array and restriction-associated DNA (RAD) sequencing. Overall, we identified 14 moderate-large effect QTL that explained up to 60.8% of the genetic variance in one of the two populations and 27.7% in the other. Four of these QTL were found in both populations explaining a substantial proportion of the variance, although major differences were also detected between the two populations. Our results confirm that BCWD resistance is controlled by the oligogenic inheritance of few moderate-large effect loci and a large-unknown number of loci each having a small effect on BCWD resistance. We detected differences in QTL number and genome location between two GWAS models (weighted single-step GBLUP and Bayes B), which highlights the utility of using different models to uncover QTL. The RAD-SNPs detected a greater number of QTL than the 57K SNP array in one population, suggesting that the RAD-SNPs may uncover polymorphisms that are more unique and informative for the specific population in which they were discovered.

## INTRODUCTION

Bacterial cold water disease (BCWD) causes significant mortality and economic losses in salmonid aquaculture (Nematollahi *et al.* 2003; Barnes and brown 2011). The etiological agent of BCWD is a gram-negative bacterium, *Flavobacterium psychrophilum (Fp)*, and current methods to control BCWD outbreaks are limited. At the National Center for Cool and Cold Water Aquaculture (NCCCWA), we have developed a selective breeding program to increase rainbow trout genetic resistance against BCWD and have shown that BCWD resistance is a moderately heritable trait that responds to selection (Leeds *et al.* 2010). Furthermore, we revealed complex genetic architecture of BCWD resistance (Vallejo *et al.* 2010) and identified several moderate-large effect quantitative trait loci (QTL) for this trait in the NCCCWA odd- and even-year rainbow trout selective-breeding populations (Wiens *et al.* 2013; Vallejo *et al.* 2014a; Liu *et al.* 2015b; Palti *et al.* 2015b). While those loci can be fine mapped to identify positional candidate genes, the complex genetic architecture of BCWD resistance and high genetic variability discovered in past studies (Vallejo *et al.* 2014a) suggest that whole genome-enabled selection is more effective for improving genetic resistance against BCWD in rainbow trout aquaculture, and we were able to empirically demonstrate that whole genome-enabled selection can double the accuracy of predicted genetic merit of potential breeders compared to traditional family-based selection (Vallejo *et al.* 2017).

For agricultural livestock species, single nucleotide polymorphism (SNP) chips have been the platform of choice for whole genome genotyping of at least 50K SNPs (Matukumalli *et al.* 2009; Ramos *et al.* 2009; Groenen *et al.* 2011); including the recently developed 57K SNP chip for rainbow trout (Palti *et al.* 2015a). However, sequencing-by-genotyping methods that do not require *a priori* marker discovery or a reference genome sequence and are capable of simultaneous marker discovery and genotyping in many individuals were developed for genetic analyses (Davey *et al.* 2011). One such technique is restriction-site-associated DNA (RAD) sequencing (Miller *et al.* 2007; Baird *et al.* 2008). In recent years, the method of RAD genotyping by sequencing has been widely used in salmonid species for SNP discovery, generating linkage maps, QTL mapping, genome-wide association studies (GWAS) and for evaluating genome-enabled selection (Hecht *et al.* 2012; Houston *et al.* 2012; Miller *et al.* 2012; Hale *et al.* 2013; Hecht *et al.* 2013; Narum *et al.* 2013; Brieuc *et al.* 2014; Campbell*et al.* 2014; Gonen *et al.* 2014; Houston *et al.* 2014; Palti *et al.* 2014; Liu *et al.* 2015a; Liu *et al.* 2015b; Palti *et al.* 2015b; Vallejo *et al.* 2016).

Several moderate-large effect QTL associated with BCWD resistance have been identified on 24 of the 29 rainbow trout chromosomes using linkage analysis mapping (Johnson *et al.* 2008; Wiens *et al.* 2013; Palti *et al.* 2014; Quillet *et al.* 2014; Vallejo *et al.* 2014a) and GWAS methods (Campbell *et al.* 2014; Liu *et al.* 2015b; Palti *et al.* 2015b). However, these previous studies had limitations on QTL detection. First, most of the reported genome-wide association analyses have tested one SNP at a time using single-regression or mixed linear models with a fixed SNP effect along with a random polygenic effect to capture the effects of all other genes. Although those studies have been successful in detecting associations, those associations typically explain only a small fraction of the trait genetic variance (Visscher *et al.* 2010). Conversely, in GWAS analysis using whole-genome selection models that simultaneously fit all SNPs as random effects, the SNPs jointly explain a larger proportion of the genetic variance which highlights the utility of using multiple-regression GWAS models with the SNPs joined into genomic windows for accurate QTL mapping (Hayes *et al.* 2010; Fan *et al.* 2011; Onteru *et al.* 2011). Second, most of the reported QTL for BCWD resistance were identified using linkage-based methods and GWAS was performed within individual segregating families with relatively small sample size and low detection power. Consequently about half of the reported findings from such QTL mapping studies are expected to represent false positives (Higginson and munafo 2016). Third, the previous studies used lower density genotyping platforms and did not have a reference genome physical map for accurate prediction of the order and physical proximity of the genetic markers. Furthermore, to our knowledge, in the current study we are using for the first time SNP genotype data with the genome physical map coordinates for GWAS in rainbow trout.

There is ambiguity on the best computational algorithm when using multiple-regression based models in genomic selection (GS) and GWAS experiments. The genetic architecture of the trait and the population structure can have a significant impact on the accuracy of the genomic predictions and estimated marker effects. Therefore, it is important to compare the performance of the best competing algorithms on GS and GWAS when evaluating a trait with complex inheritance for the first time in a population. This will allow effective discovery of QTL underlying the genetic basis of the complex trait and control the type I error rate, which is often high in genome-wide discovery experiments.

In GWAS with models that fit all SNPs simultaneously, the genomic BLUP (GBLUP) method assumes a polygenic architecture of the trait and uses all SNP data to estimate the genomic relationship (G) matrix. In contrast, the Bayesian variable selection method assumes that the genetic variance is explained by a reduced number of markers with small-moderate or large effects (Habier *et al.* 2007; Hayes *et al.* 2009; De los campos *et al.* 2013; Fernando and garrick 2013; Howard *et al.* 2015). Based on this assumption, GBLUP is expected not to perform as well as Bayesian variable selection models when the trait is controlled by few moderate-to-large effect QTL. The GBLUP method was modified into the single-step GBLUP (ssGBLUP) method which allows combining the pedigree (A) and genomic-derived relationships into an H relationship matrix (Aguilar *et al.* 2010; Legarra *et al.* 2014), and to the weighted single-step GBLUP (wssGBLUP) method which emulates the Bayesian variable selection models by fitting in the multiple regression model selected SNPs that explain moderate-large fraction of the trait genetic variation (Wang *et al.* 2012).

The recent development of the 57K SNP array (Palti *et al.* 2015a), a dense genetic linkage map with 47,939 SNP markers (Gonzalez-pena *et al.* 2016), and the release of the improved rainbow trout reference genome (GenBank assembly Accession GCA_002163495) have provided the needed tools for performing GWAS to identify genomic regions associated with BCWD resistance in rainbow trout. The main objectives of this study were to (1) identify and validate QTL associated with BCWD resistance in two commercially-relevant rainbow trout breeding populations; (2) characterize the genetic architecture of rainbow trout resistance to BCWD; (3) compare the QTL mapping efficiency and determine whether the Chip-SNP and RAD-SNP genotyping platforms detect the same QTL; and (4) compare the QTL mapping efficiency of two widely-used multiple-regression GWAS models.

## MATERIALS AND METHODS

### Ethics statement

Protocols for this study were reviewed and approved by the NCCCWA Institutional Animal Care and Use Committee (Kearneysville, WV).

### Rainbow trout rearing and BCWD challenge

Details of the 21-day BCWD challenge have been reported elsewhere (Silverstein *et al.* 2009; Leeds *et al.* 2010). Mortalities were removed and recorded daily and fin clipped. Fish that survived the challenge were euthanized in a lethal dose of Tricaine methane sulfonate (Tricaine-S, Western Chemical, Inc., Ferndale, WA) and fin clipped. Fin clips from all fish (mortalities and survivors) were individually kept in 95% ethanol until DNA was extracted using established protocols (Palti *et al.* 2006).

### Rainbow trout populations used in GWAS

Fish used in this study were sampled from two populations, and all analyses were performed separately by population. The first sample included fish with genotypes and phenotypes from 10 full-sib (FS) families sampled from a total of 71 pedigreed FS families with phenotype data from year-class (YC) 2005 of the NCCCWA BCWD resistant line (NCCCWA) and it was described in our previous GS study (Vallejo et al., 2016). Briefly, the YC 2005 families represented the base generation of the breeding line, and thus had not previously been selected for BCWD resistance. Each family had *n* = 39-80 fish evaluated in the laboratory BCWD challenge in one or two tanks per family. The phenotypic dataset included disease resistance phenotypes from *n* =4,492 fish from 71 FS families (Table 1), and the pedigree file included 4,659 records. In this NCCWA sample, a total of *n* = 583 fish had both genotype and phenotype records. Following pedigree quality control, the original NCCCWA sample of *n* = 583 genotyped fish was reduced into *n* = 577 because four fish were flagged as duplicated or cloned samples by the QC pipeline and two fish did not assign to the expected family based on the pedigree records.

**Table 1.**
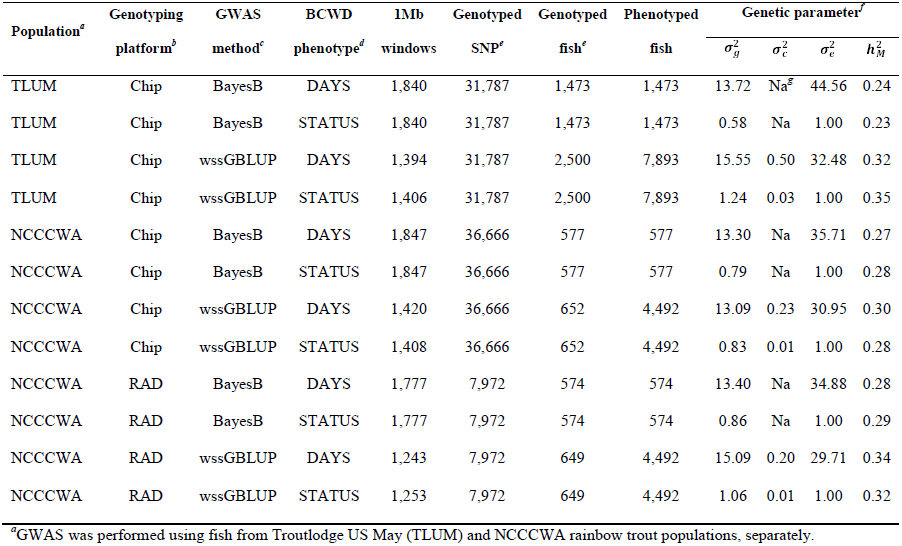
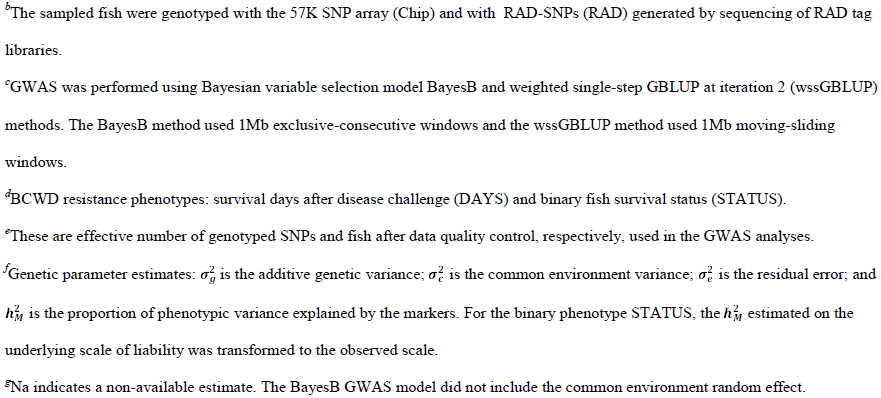
Experimental variables of GWAS conducted in two rainbow trout populations using two SNP genotyping methods.

| Population<sup>a</sup> | Genotyping platform<sup>b</sup> | GWAS method<sup>c</sup> | BCWD phenotype<sup>d</sup> | 1Mb windows | Genotyped SNP<sup>e</sup> | Genotyped fish<sup>e</sup> | Phenotyped fish | Genetic parameter<sup>f</sup> |  |  |  |
| --- | --- | --- | --- | --- | --- | --- | --- | --- | --- | --- | --- |
| | | | | | | | | $\sigma_g^2$ | $\sigma_c^2$ | $\sigma_e^2$ | $h_M^2$ |
| TLUM | Chip | BayesB | DAYS | 1,840 | 31,787 | 1,473 | 1,473 | 13.72 | Na<sup>g</sup> | 44.56 | 0.24 |
| TLUM | Chip | BayesB | STATUS | 1,840 | 31,787 | 1,473 | 1,473 | 0.58 | Na | 1.00 | 0.23 |
| TLUM | Chip | wssGBLUP | DAYS | 1,394 | 31,787 | 2,500 | 7,893 | 15.55 | 0.50 | 32.48 | 0.32 |
| TLUM | Chip | wssGBLUP | STATUS | 1,406 | 31,787 | 2,500 | 7,893 | 1.24 | 0.03 | 1.00 | 0.35 |
| NCCCWA | Chip | BayesB | DAYS | 1,847 | 36,666 | 577 | 577 | 13.30 | Na | 35.71 | 0.27 |
| NCCCWA | Chip | BayesB | STATUS | 1,847 | 36,666 | 577 | 577 | 0.79 | Na | 1.00 | 0.28 |
| NCCCWA | Chip | wssGBLUP | DAYS | 1,420 | 36,666 | 652 | 4,492 | 13.09 | 0.23 | 30.95 | 0.30 |
| NCCCWA | Chip | wssGBLUP | STATUS | 1,408 | 36,666 | 652 | 4,492 | 0.83 | 0.01 | 1.00 | 0.28 |
| NCCCWA | RAD | BayesB | DAYS | 1,777 | 7,972 | 574 | 574 | 13.40 | Na | 34.88 | 0.28 |
| NCCCWA | RAD | BayesB | STATUS | 1,777 | 7,972 | 574 | 574 | 0.86 | Na | 1.00 | 0.29 |
| NCCCWA | RAD | wssGBLUP | DAYS | 1,243 | 7,972 | 649 | 4,492 | 15.09 | 0.20 | 29.71 | 0.34 |
| NCCCWA | RAD | wssGBLUP | STATUS | 1,253 | 7,972 | 649 | 4,492 | 1.06 | 0.01 | 1.00 | 0.32 |

<sup>a</sup>GWAS was performed using fish from Troutlodge US May (TLUM) and NCCCWA rainbow trout populations, separately.
<sup>b</sup>The sampled fish were genotyped with the 57K SNP array (Chip) and with RAD-SNPs (RAD) generated by sequencing of RAD tag libraries.
<sup>c</sup>GWAS was performed using Bayesian variable selection model BayesB and weighted single-step GBLUP at iteration 2 (wssGBLUP) methods. The BayesB method used 1Mb exclusive-consecutive windows and the wssGBLUP method used 1Mb moving-sliding windows.
<sup>d</sup>BCWD resistance phenotypes: survival days after disease challenge (DAYS) and binary fish survival status (STATUS).
<sup>e</sup>These are effective number of genotyped SNPs and fish after data quality control, respectively, used in the GWAS analyses.
<sup>f</sup>Genetic parameter estimates: $\sigma_g^2$ is the additive genetic variance; $\sigma_e^2$ is the common environment variance; $\sigma_e^2$ is the residual error; and $h_M^2$ is the proportion of phenotypic variance explained by the markers. For the binary phenotype STATUS, the $h_M^2$ estimated on the underlying scale of liability was transformed to the observed scale.
<sup>g</sup>Na indicates a non-available estimate. The BayesB GWAS model did not include the common environment random effect.

The second sample included 102 pedigreed FS families from YC 2013 of the commercial breeding company Troutlodge, Inc., May-spawning population (TLUM) and it was described in our previous GS study (Vallejo *et al.* 2017). Briefly, the original study design was to sample *n* = 1500 fish with phenotypes and genotypes from 50 FS families; in practice, a total of *n* = 1473 fish had both phenotype and genotype records from those 50 FS families (*n* = 17-40 fish per family). The 102 YC 2013 families represented a commercial nucleus breeding population undergoing selection for growth, and thus had not previously been selected for BCWD resistance. The fish were evaluated for BCWD survival in the laboratory challenge in two tanks per family with an initial stocking of 40 fish per tank. The phenotypic dataset included BCWD disease phenotype records from *n* = 7893 fish from 102 FS families (Table 1), and the pedigree file included 32,279 records. A summary of the experimental variables of GWAS conducted with fish sampled from these two rainbow trout populations is presented in Table 1.

### BCWD resistance phenotypes

The discrete BCWD resistance phenotype DAYS, the number of days post-challenge until the fish succumbed to the disease, was recorded for all mortalities and survivors were assigned a value of 21. Each fish also had a binary survival STATUS record. The BCWD resistance phenotype STATUS had two categories: 1 = the fish died during the 21 days post challenge evaluation period; and 2 = the fish survived for the duration of the challenge. The DAYS and STATUS records were analyzed separately using univariate GWAS models described below.

### SNP genotyping platforms

The fish sampled from TLUM and NCCCWA populations were genotyped using the Rainbow Trout Affymetrix 57K SNP array (Chip) following previously described procedures (Palti *et al.* 2015a) and the samples were genotyped by a commercial service provider (Geneseek, Inc., Lincoln, NE) following the Axiom genotyping procedures described by the array manufacturer (Affymetrix). The quality control (QC) bioinformatics pipeline applied to the Chip-SNP genotype data collected in the TLUM (Vallejo *et al.* 2017) and NCCCWA (Vallejo *et al.* 2016) populations were already described. After genotype data QC, a total of 41,868 and 49,468 SNPs were included in the TLUM and NCCCWA raw Chip genotype datasets, respectively.

The fish sampled from the NCCCWA population were also genotyped by sequencing of restriction-site-associated DNA (RAD) tag libraries as we have previously described (Vallejo *et al.* 2016). After genotype data QC, a total of 24,465 RAD-SNPs were included in the raw RAD genotype dataset. The raw sequence data from the RAD libraries were deposited in the NCBI SRA database (Accession SRP063932).

Before performing GWAS analyses, the raw marker genotype datasets were further QC filtered using algorithms implemented in the software BLUPF90 (Misztal *et al.* 2015). For the Chip data, the QC retained SNPs with a genotype calling rate higher than 0.90, minor allele frequency higher than 0.05, and departures from Hardy-Weinberg equilibrium less than 0.15 (difference between expected and observed frequency of heterozygotes). Parent-progeny pairs were tested for discrepant homozygous SNPs, and SNPs with a conflict rate of more than 1% were discarded from further analysis. For the RAD data, the QC retained SNPs with a genotype calling rate higher than 0.70. Following this final QC step, 33,838 SNPs, 39,112 SNPs and 9,534 SNPs were retained for analyses of the TLUM, NCCCWA (Chip) and NCCCWA (RAD) datasets, respectively. Next, we determined the physical map position (GenBank assembly Accession GCA_002163495) of each of the QC filtered markers and found that a small fraction did not have a physical map location. The numbers of effective genotyped markers and effective genotyped fish that were used with each specific GWAS model and genotyping platform in the evaluated populations are shown in Table 1.

### Genome-wide association analyses

This study was conducted to identify chromosomal regions that have the greatest impact on the variation of BCWD resistance because it is difficult to infer individual SNP effects from a multiple regression model that fits markers simultaneously at a genome-wide scale (Garrick and fernando 2013). Therefore, instead of using the markers effect to make an inference on a particular locus, we used the markers effect to make an inference about a particular genomic region that encompasses a number of contiguous loci in association with the trait (Fan *et al.* 2011; Fernando *et al.* 2014). Thus the GWAS analysis was performed separately for each population and BCWD resistance phenotype using multiple-regression GWAS models that use simultaneously all the SNPs in the association test. Two multiple-regression GWAS methods were used: Bayesian variable selection BayesB (BayesB) (Fernando and garrick 2009) and weighted single-step GBLUP (wssGBLUP) (Wang *et al.* 2012; Misztal *et al.* 2015).

Before starting the search for genomic regions associated with BCWD resistance, we tested 0.5 and 1Mb exclusive-windows in GWAS with BayesB model and found that 1Mb windows provided Manhattan plots with less noisy baseline which was in agreement with other GWAS reports (Kizilkaya *et al.* 2013; Saatchi *et al.* 2013); so we decided to use 1Mb exclusive-windows in the GWAS performed with BayesB. We also used 1Mb sliding-windows in GWAS performed with wssGBLUP which are more informative than exclusive-windows.

### Bayesian variable selection model BayesB

The BayesB model uses only fish that had both genotype and phenotype records. The TLUM population sample included *n* = 1473 fish from 50 YC 2013 families with phenotype and genotype records, and the NCCWA sample included *n* = 577 fish from 10 YC 2005 families with phenotype and genotype records (Table 1).

The GWAS for DAYS was performed using this linear model: y = 1*μ*+ Z*α* + *e*; where y is *n x 1* vector of phenotypic records; **1** is a vector of all ones; μ is overall mean of phenotypic records; Z is an *n x k* matrix of genotype covariates (coded as -10, 0, or 10) for *k* SNP markers, *α* is a *k x 1* vector of random regression coefficients of ***k*** additive marker effects, and ***e*** is a vector of residuals. The genotype and phenotype records were used to estimate markers effect using the Bayesian variable selection model BayesB implemented in the software GENSEL (Fernando and garrick 2009) as we have previously described (Vallejo *et al.* 2016). The GWAS for the binary phenotype STATUS was performed using the option for categorical analysis implemented also in the software GENSEL (Fernando and garrick 2009; Fernando and garrick 2013; Garrick and fernando 2013).

The BayesB model fits a mixture model to estimate marker effects assuming there are two types of SNP markers: a fraction of non-zero SNP effects (1-π) that are drawn from distributions with marker-specific variance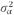, and other known fraction of markers (π) that have zero effect on the quantitative trait (Meuwissen *et al.* 2001). In this study, the mixture parameter π was determined empirically by testing each population dataset listed in Table 1 (TLUM-Chip, NCCCWA-Chip and NCCCWA-RAD) with several π values (0.95, 0.96, 0.97, 0.98 and 0.99). We then decided to use π values of 0.97 and 0.99 with TLUM-Chip and NCCCWA samples, respectively, in the GWAS analysis with BayesB because these π values yielded best accuracy genomic predictions (Results not presented).

The BayesB model uses Gibbs sampling approach in the GWAS analysis (Garrick and fernando 2013). In this study, the BCWD phenotypes were analyzed using 270,000 Markov Chain Monte Carlo (MCMC) iterations from which the first 20,000 samples were discarded as burn-in; from the remaining 250,000 samples, we saved one from every 50 samples so the marker effects and variances were estimated as the posterior means of collected 5,000 independent samples. The proper mixing and convergence of the MCMC iterations were assessed with the R package CODA (Plummer *et al.* 2006).

### Weighted single-step GBLUP model

The GWAS analysis with wssGBLUP, in contrast to BayesB, uses all available information on sampled fish such as pedigree, genotype and phenotype records including those fish that had only phenotypic records (i.e., those with missing genotype data) as long as the sampled fish are pedigree related (Aguilar *et al.* 2010; Christensen AND LUND 2010). The TLUM sample included *n* = 7893 fish from 102 YC 2013 families with phenotype records, and the NCCWA sample included *n* = 4492 BCWD fish from 71 YC 2005 families with phenotype records (Table 1).

In GWAS with wssGBLUP, the weights for each SNP are 1’s for the 1^st^ iteration which means that all SNPs have the same weight (i.e., standard single-step GBLUP). For the next iterations (2^nd^, 3^rd^, etc.), the weights are SNP specific variances calculated using both the SNP allele-substitution effect estimated in the previous iteration and their corresponding allele frequencies (Wang *et al.* 2012). In this study, we decided to use results from the 2^nd^ iteration because they provide the highest accuracy genomic predictions (Vallejo *et al.* 2016) and marker effects (Wang *et al.* 2012; Irano *et al.* 2016; Melo *et al.* 2016).

In GWAS with TLUM population sample, the linear and threshold models for DAYS and STATUS, respectively, included the effects of population mean, random animal, random common environment and random error. The full-sib fish progeny from each family was allocated into two tanks for BCWD challenge evaluation, so the variable tank nested within family was used to model the common environment effect. The linear and threshold models used with the NCCCWA sample were similar to those used with the TLUM sample. The linear model for DAYS and threshold model for STATUS were fitted using computer applications implemented in the software BLUPF90 (Misztal *et al.* 2015). The binary phenotype disease STATUS was analyzed with a threshold model using a Bayesian approach which included a single chain with a total of 270,000 iterations; the first 20,000 iterations were discarded as burn-in iterations; then from the remaining 250,000 samples, one from every 50 samples were saved. Thus, 5000 independent samples were used in the analysis. The proper mixing and convergence of these MCMC iterations were also assessed using the R package CODA (Plummer *et al.*2006).

### Criteria to declare QTL associated with BCWD resistance

The results from the GWAS performed with BayesB and wssGBLUP were used to identify genomic windows and QTL associated with BCWD resistance. A two stage approach was used to identify a QTL associated with BCWD resistance. First, the genomic windows with explained genetic variance (EGV) greater than 1% and 2% in the TLUM and NCCCWA populations, respectively, were declared as genomic regions associated with BCWD resistance. The threshold to declare a QTL was raised to EGV≥2% in the relatively small NCCCWA sample (*n* = 577) to control the type I error rate. Second, to determine if neighboring or overlapping windows on the same chromosome belong to the same QTL region we used the following criteria: all windows associated with BCWD resistance that were bounded within a region smaller than 20Mb and were less than 10Mb apart from another associated window were grouped to a single QTL region. The QTL nomenclature we used was based on the chromosome number and QTL region within the chromosome, where the region with the lowest genome assembly position numbers determined to be QTL1 on that chromosome the next QTL2 and so on. For example, the QTL regions detected on chromosome Omy3 and listed in Table 2 were separated to QTL 3.1, 3.2 and 3.3 based on the physical genome map positions of the SNPs that flanked each QTL region.

**Table 2.** Summary of QTL associated with BCWD survival STATUS in the Troutlodge US May (TLUM) population^*a*^.

| Omy | QTL<sup>b</sup> | GWAS<br>method<sup>c</sup> | Genetic<br>variance<br>(%)<sup>d</sup> | Physical Map (bp)<sup>e</sup> |  | Markers in Window |  | SNPs<br>per<br>window |
| --- | --- | --- | --- | --- | --- | --- | --- | --- |
|  |  |  |  | Start | End | Start-SNP | End-SNP |  |
| 3 | 3.1 | wssGBLUP | 1.8 | 17,812,341 | 18,600,963 | Affx-88909970 | Affx-88904917 | 22 |
| 3 | 3.2 | wssGBLUP | 2.0 | 61,621,949 | 62,558,467 | Affx-88925949 | Affx-88919479 | 18 |
| 3 | 3.3 | wssGBLUP | 1.2 | 77,108,538 | 78,076,592 | Affx-88925305 | Affx-88929879 | 19 |
| 5 | 5.1 | wssGBLUP | 1.2 | 11,339,155 | 12,329,117 | Affx-88916119 | Affx-88936955 | 27 |
| 8 | 8.1 | BayesB | 19.3 | 76,070,399 | 76,907,400 | Affx-88955037 | Affx-88906927 | 17 |
| 25 | 25.1 | BayesB | 35.4 | 21,006,787 | 21,805,909 | Affx-88924154 | Affx-88936445 | 30 |

<sup>a</sup>The fish from TLUM population were genotyped with the 57K SNP array (Chip).
<sup>b</sup>From each QTL, the window with the highest explained genetic variance is presented in this Table.
<sup>c</sup>GWAS was conducted using Bayesian variable selection model BayesB (BayesB) and weighted single-step GBLUP (wssGBLUP) methods. BayesB used 1Mb exclusive-consecutive windows and wssGBLUP used 1Mb moving-sliding windows.
<sup>d</sup>Explained genetic variance by tested window (%).
<sup>e</sup>SNP positions in base pairs (bp) based on rainbow trout reference genome sequence (GenBank assembly Accession GCA\_002163495).

The BayesB algorithm estimated the proportion of models where the tested 1Mb-window was included as with non-zero variance (p > 0) and therefore accounted for more than 0% of the genetic variance. These p > 0 estimates were used to calculate the probability of false positives [PFP = 1 - (p > 0)] for each tested window (Fernando and garrick 2013). These PFP estimates enabled accounting for multiple testing to control the probability of false positive conclusions across all the undertaken GWAS tests with the BayesB model. Thus by using these EGV thresholds (Peters *et al.* 2013) and PFP estimates (Fernando and garrick 2013), the claims on significant QTL findings was restricted to the strongest associations with BCWD resistance to control the type I error rate in this study.

### QTL segregating in both NCCCWA and TLUM populations

In order to identify QTL that might be overlapping between the two populations and between the two genotyping platforms in the NCCCWA population, we assigned the genome physical map positions to all flanking markers of the genomic windows associated with BCWD resistance using the rainbow trout reference genome sequence (GenBank assembly Accession GCA_002163495) and searched for overlapping QTL regions within each chromosome using the flanking markers physical map genome coordinates.

### Data availability

The authors state that all data necessary for supporting the conclusions of this research article are included within the article and it’s Supplemental Material. Figure S1 shows Manhattan plot with GWAS results for survival DAYS in TLUM sample genotyped with 57K Chip-SNP. Figure S2 shows Manhattan plot with GWAS results for survival DAYS in NCCCWA sample genotyped with 57K Chip-SNP. Figure S3 shows Manhattan plot with GWAS results for survival DAYS in NCCCWA sample genotyped with RAD-SNPs. Table S1 contains summary of all genomic windows and QTL detected in this GWAS. Table S2 contains list of QTL associated with survival DAYS found in TLUM population using the 57K Chip-SNP. Table S3 contains list of QTL associated with survival DAYS found in NCCCWA population using the 57K Chip-SNP. Table S4 contains list of QTL associated with survival DAYS found in NCCCWA population using RAD-SNPs genotyping. Table S5 contains list of QTL that are segregating in both NCCCWA and TLUM populations. Table S6 contains list of QTL that are private to either NCCCWA or TLUM population. Table S7 contains summary of QTL for BCWD resistance reported in previous studies. The rainbow trout reference genome sequence is available at GenBank with assembly accession number: GCA_002163495.

## RESULTS

### Heritability of BCWD resistance

The heritability or proportion of phenotypic variance explained by the markers for survival DAYS and the binary survival STATUS were previously reported (Vallejo *et al.* 2016; Vallejo *et al.* 2017). Briefly, here they were moderate and relatively constant with a range of 0.24-0.34 (DAYS) and 0.23-0.35 (STATUS) (Table 1); and the mean heritability for DAYS and STATUS were similar (0.29). The heritability of STATUS estimated on the underlying scale of liability using a threshold model was transformed to the observed scale of disease survival STATUS using already described procedures (Vallejo *et al.* 2017). Overall, for both BCWD phenotypes and across genotyping platforms and populations, the mean heritability estimated with wssGBLUP (0.32) was slightly higher than that estimated with BayesB (0.27).

### QTL associated with BCWD resistance in the TLUM population (57K SNP Chip)

We have previously shown that the BCWD resistance phenotypes DAYS and STATUS yielded similar results in QTL mapping (Vallejo *et al.* 2014a; Liu *et al.* 2015b; Palti *et al.* 2015b) and genomic selection experiments (Vallejo *et al.* 2016; Vallejo *et al.* 2017). In this study, we observed also that STATUS and DAYS were affected by similar QTL regions with few exceptions (Overall, DAYS detected three more QTL than STATUS) (Table S1). However, because STATUS resembles the disease resistance trait closer than DAYS; and selection programs for improved disease resistance would most likely favor improvement of resistance over endurance or tolerance (Odegard *et al.* 2011), we present the results from the STATUS survival phenotype in the main body of this report. The complete results from the analysis with the DAYS phenotype are presented in the Supplementary Material section.

In the TLUM population, a total of 45 windows with EGV ≥1% were detected on chromosomes Omy3, 5, 8, 13 and 25 (Table 2; Figure 1; Table S1 and Figure S1). Fourteen windows were detected with BayesB with EGV up to 57.6% and 31 windows were detected with wssGBLUP with EGV up to 28.7%.

**Figure 1.**
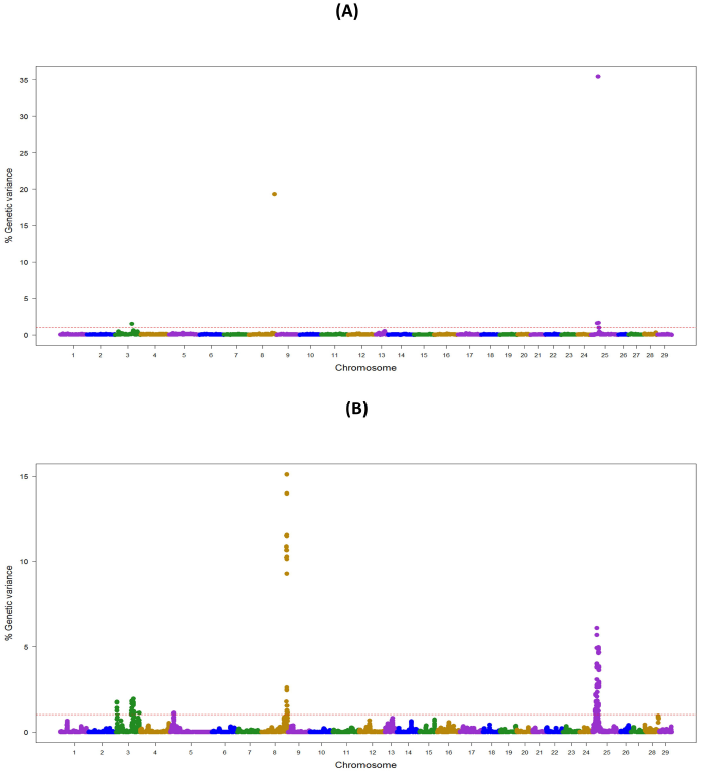
Manhattan plot showing the association between SNP genomic windows and BCWD resistance in TLUM sample genotyped with 57K Chip-SNP: (A) GWAS for STATUS performed with BayesB using 1Mb exclusive windows. (B) GWAS for STATUS performed with wssGBLUP using 1Mb sliding windows. (PPTX file)

Four QTL (3.2, 8.1, 13.2 and 25.1) were detected by both GWAS models (Table S1). We did not detect any BayesB model specific QTL (i.e., all QTL detected with BayesB were also detected with wssGBLUP); and four QTL (3.1, 3.3, 5.1 and 13.1) were detected only with wssGBLUP.

Overall, we detected eight QTL (3.1, 3.2, 3.3, 5.1, 8.1, 13.1, 13.2 and 25.1) associated with BCWD resistance which jointly explained up to 61% of the genetic variance for BCWD resistance in this TLUM population when accounting only for the largest EGV window in each QTL (Table 2; Table S1 and S2). Among these eight QTL, two significant large-effect QTL were detected on Omy8 (QTL 8.1; *PFP*= 0.0) and 25 (QTL 25.1; *PFP*= 0.01) with BayesB, each explaining up to 19.3% and 35.4% of the genetic variance for BCWD resistance, respectively (Table 2).

### QTL for BCWD resistance in the NCCCWA population (57K SNP Chip)

In the NCCWA population using the 57K SNP chip, we detected a total of 11 windows associated with BCWD resistance on chromosomes Omy3, 5, 10, 22 and 25 (Table 3; Figure 2; Table S1 and Figure S2). Two and nine QTL windows were detected with BayesB and wssGBLUP, respectively, and each GWAS model explained up to 5.6% and 16.1% of the genetic variance, respectively.

**Table 3.**
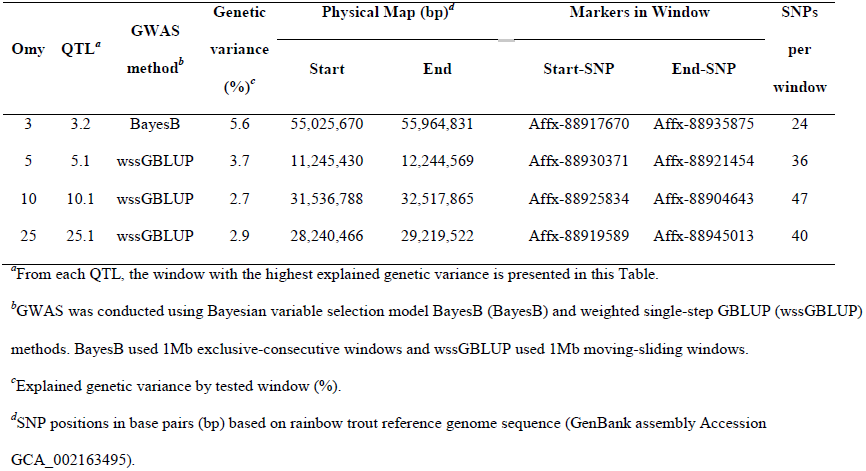
Summary of QTL associated with BCWD survival STATUS in NCCCWA population detected using the 57K SNP.

| Omy | QTL<sup>a</sup> | GWAS<br>method<sup>b</sup> | Genetic<br>variance<br>(%)<sup>c</sup> | Physical Map (bp)<sup>d</sup> |  | Markers in Window |  | SNPs<br>per<br>window |
| --- | --- | --- | --- | --- | --- | --- | --- | --- |
|  |  |  |  | Start | End | Start-SNP | End-SNP |  |
| 3 | 3.2 | BayesB | 5.6 | 55,025,670 | 55,964,831 | Affx-88917670 | Affx-88935875 | 24 |
| 5 | 5.1 | wssGBLUP | 3.7 | 11,245,430 | 12,244,569 | Affx-88930371 | Affx-88921454 | 36 |
| 10 | 10.1 | wssGBLUP | 2.7 | 31,536,788 | 32,517,865 | Affx-88925834 | Affx-88904643 | 47 |
| 25 | 25.1 | wssGBLUP | 2.9 | 28,240,466 | 29,219,522 | Affx-88919589 | Affx-88945013 | 40 |

<sup>a</sup>From each QTL, the window with the highest explained genetic variance is presented in this Table.
<sup>b</sup>GWAS was conducted using Bayesian variable selection model BayesB (BayesB) and weighted single-step GBLUP (wssGBLUP) methods. BayesB used 1Mb exclusive-consecutive windows and wssGBLUP used 1Mb moving-sliding windows.
<sup>c</sup>Explained genetic variance by tested window (%).
<sup>d</sup>SNP positions in base pairs (bp) based on rainbow trout reference genome sequence (GenBank assembly Accession GCA\_002163495).

**Figure 2.**
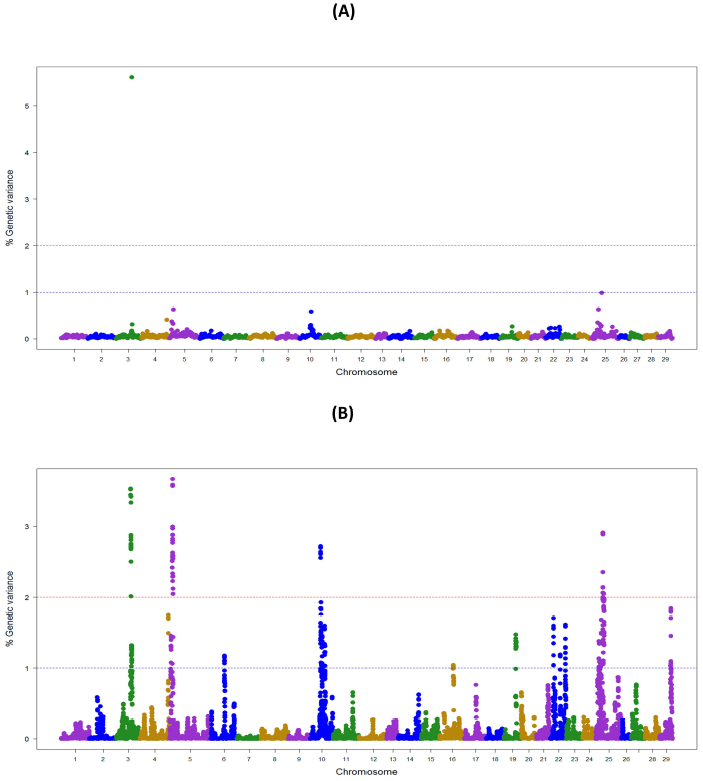
Manhattan plot showing the association between SNP genomic windows and BCWD resistance in NCCCWA sample genotyped with 57K Chip-SNP: (A) GWAS for STATUS performed with BayesB using 1Mb exclusive windows. (B) GWAS for STATUS performed with wssGBLUP using 1Mb sliding windows. (PPTX file)

Only one QTL (3.2) was detected by both GWAS models (Table S1). We did not detect any BayesB model specific QTL, and four QTL (5.1, 10.1, 22.1 and 25.1) were detected only with the wssGBLUP method.

Overall, we detected five QTL (3.2, 5.1, 10.1, 22.1 and 25.1) associated with BCWD resistance that explained up to 18.2% of the genetic variance in the NCCCWA population when accounting only for the largest EGV window in each QTL (Table 3 and Table S3).

### QTL for BCWD resistance in the NCCCWA population detected with RAD SNPs

In the NCCCWA population using the RAD SNP genotypes, we detected a total of 18 windows associated with BCWD resistance on chromosomes Omy3, 5, 10, 11, 13, 15 and 25 (Table 4; Figure 3; Table S1 and Figure S3). Four windows were detected by BayesB with EGV up to 17.3% and 14 windows were detected by wssGBLUP with EGV up to 26.4%.

**Table 4.** Summary of QTL associated with BCWD survival STATUS in NCCCWA population detected using RAD-SNPs.

| Omy | QTL<sup>a</sup> | GWAS method<sup>b</sup> | Genetic variance (%)<sup>c</sup> | Physical Map (bp)<sup>d</sup> |  | Markers in Window |  | SNPs per window |
| --- | --- | --- | --- | --- | --- | --- | --- | --- |
|  |  |  |  | Start | End | Start-SNP | End-SNP |  |
| 3 | 3.2 | BayesB | 5.1 | 55,254,048 | 55,993,431 | BCWD10F04977 | BCWD10F00765 | 3 |
| 5 | 5.1 | wssGBLUP | 2.8 | 11,966,155 | 12,948,479 | BCWD10F15578 | BCWD10F00357 | 11 |
| 5 | 5.2 | BayesB | 3.3 | 41,094,726 | 41,686,801 | BCWD10F02753 | BCWD10F18861 | 6 |
| 13 | 13.1 | wssGBLUP | 2.1 | 11,230,016 | 11,602,309 | BCWD10F05067 | BCWD10F24318 | 3 |
| 15 | 15.1 | wssGBLUP | 3.7 | 38,446,758 | 39,349,228 | BCWD10F00773 | BCWD10F16617 | 3 |
| 25 | 25.1 | wssGBLUP | 6.6 | 17,496,495 | 18,444,865 | BCWD10F06483 | BCWD10F14339 | 11 |

<sup>a</sup>From each QTL, the window with the highest explained genetic variance is presented in this Table.
<sup>b</sup>GWAS was conducted using Bayesian variable selection model BayesB (BayesB) and weighted single-step GBLUP (wssGBLUP) methods. BayesB used 1Mb exclusive-consecutive windows and wssGBLUP used 1Mb moving-sliding windows.
<sup>c</sup>Explained genetic variance by tested window (%).
<sup>d</sup>SNP positions in base pairs (bp) based on rainbow trout reference genome sequence (GenBank assembly Accession GCA\_002163495).

**Figure 3.**
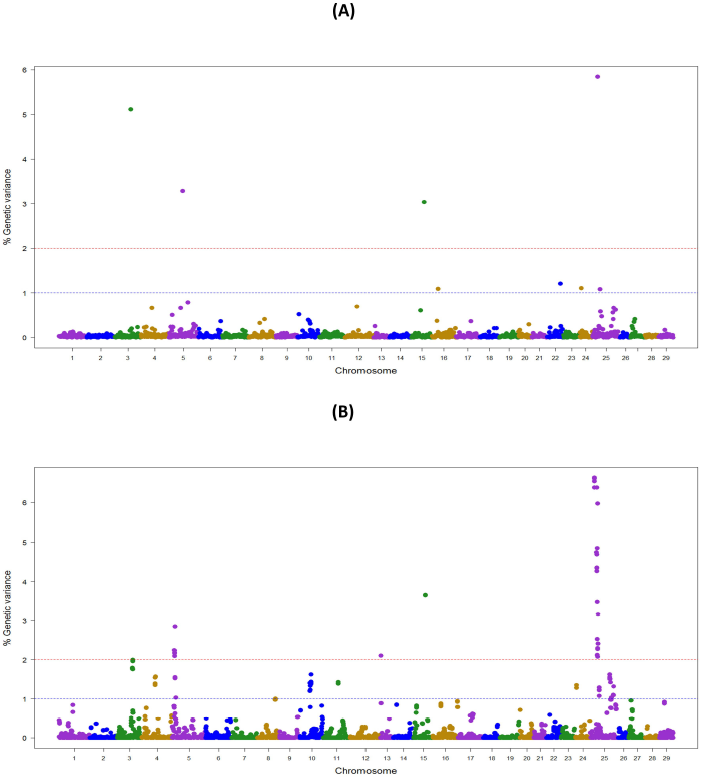
Manhattan plot showing the association between SNP genomic windows and BCWD resistance in NCCCWA sample genotyped with the RAD-SNPs: (A) GWAS for STATUS performed with BayesB using 1Mb exclusive windows. (B) GWAS for STATUS performed with wssGBLUP using 1Mb sliding windows. (PPTX file)

Three QTL (3.2, 15.1 and 25.1) were detected by both GWAS models (Table S1). QTL 5.2 was detected only with BayesB, and five QTL (5.1, 10.1, 11.1, 13.1 and 25.2) were detected only with wssGBLUP.

Overall, we detected nine QTL (3.2, 5.1, 5.2, 10.1, 11.1, 13.1, 15.1, 25.1 and 25.2) associated with BCWD resistance which explained up to 31.9% of the genetic variance in this NCCCWA population dataset when accounting only for the largest EGV window in each QTL (Table 4 and Table S4).

## DISCUSSION

In GWAS studies, the use of correct statistical models and computer algorithms is paramount to successfully identify the underlying genetic basis of resistance to complex diseases in livestock and aquaculture species. To date there have been several reported GWAS using single-marker association tests in fin fish species (Campbell *et al.* 2014; Ayllon *et al.* 2015; Geng *et al.* 2015; Gonen *et al.* 2015; Liu *et al.* 2015b; Palti *et al.* 2015b; TSai *et al.* 2015; TSai *et al.* 2016) which generally are associated with high type I error rate because single-marker methods do not account for linkage disequilibrium (LD) between physically linked loci in the association test. In this study, we performed GWAS for loci associated with BCWD resistance using multiple-regression models which estimate the effect of all markers simultaneously and consequently do account for LD between neighboring SNPs (Fernando and garrick 2013; Garrick and fernando 2013; Misztal *et al.* 2014).

In the current GWAS, we detected 14 QTL associated with BCWD resistance in two commercially-relevant rainbow trout breeding populations from which 11 were validated QTL from previous studies and three were novel QTL. Here we confirmed that BCWD resistance is controlled by the oligogenic inheritance of few moderate-large effect loci and a large-unknown number of loci with small effects on BCWD resistance. However, despite the similar genetic architecture for this trait in both populations and the detection of overlapping QTL, we still detected major QTL differences between the two populations. We also found that the RAD and Chip genotyping platforms did not detect the same QTL in the NCCCWA population, and overall the RAD platform detected a greater number of QTL than the Chip platform. In addition, the wssGBLUP and the BayesB multiple-regression GWAS models did not detect the same QTL, which highlights the utility of using different GWAS models to effectively optimize the discovery of QTL.

### The genetic architecture of BCWD resistance in rainbow trout

Previously, we predicted that six to 10 QTL explaining 83% to 89% of phenotypic variance with either additive or dominant disease-resistant alleles plus polygenic effects may underlie the genetic architecture of BCWD resistance in the same NCCCWA population using Bayesian complex segregation analysis of phenotype and pedigree records (Vallejo *et al.* 2010). In the current study we were able to confirm our prediction on the genetic architecture of the trait. With 10 families from that original NCCCWA population, we uncovered 10 moderate-large effect QTL that explained up to 27.7% of the additive genetic variance for BCWD resistance (Table S1). Similarly, in the TLUM odd-year population, we detected 8 moderate-large effect QTL that explained up to 60.8% of the additive genetic variance for BCWD resistance.

Four QTL regions located on chromosomes Omy3, 5, 13 and 25 are segregating in both populations (Table S5). The shared QTL regions explain a substantial proportion of the additive genetic variance for BCWD resistance in the two populations (up to 18% and 38.6% of the genetic variance in NCCCWA and TLUM, respectively); suggesting a common underlying genetic architecture for BCWD resistance in the two populations. However, major differences were also detected between the two populations. Six QTL, which explained up to 9.7% of the genetic variance and are located on chromosomes Omy5, 10, 11, 15, 22 and 25 were found only in the NCCCWA population (Table S6). Conversely, four QTL which explained up to 22.2% of the genetic variance and are located on chromosomes Omy3, 8 and 13 were only found in the TLUM population. Overall, our GWAS results confirmed the hypothesis that BCWD resistance is controlled by the oligogenic inheritance of several moderate-large effect QTL and many small effect polygenic loci (Vallejo *et al.* 2010; Vallejo *et al.* 2014a; Liu *et al.* 2015b; Palti *et al.* 2015b).

Further fine-mapping of the BCWD-QTL position and eventual identification of putative candidate genes or disease-causal mutations would be advantageous for applying marker-based selection and advancing the understanding of the mechanisms of genetic resistance to BCWD in rainbow trout populations. This can be achieved by genotyping and disease testing a greater number of SNPs from positions within and near the major QTL regions, by re-sequencing highly characterized BCWD resistant and susceptible individuals as was successfully done in the search for the IPNV resistance gene in Atlantic salmon (Moen *et al.* 2015). In addition, positional and functional candidate genes for the QTL can be generated by interrogating the newest version of the rainbow trout reference genome sequence (GenBank assembly Accession GCA_002163495).

### Comparing the two SNP genotyping technologies in the NCCCWA population

Overall, the RAD genotyping technology (18 windows; EGV= 32.8%; Table S1) detected a greater number of windows associated with BCWD resistance than the Chip technology (11 windows; EGV= 18.2%) in the NCCCWA population. From the 10 QTL found in the NCCCWA population, more than half of the detected QTL were genotyping platform specific: One QTL was detected only with the Chip technology (QTL 22.1); and five QTL were detected only with the RAD technology (QTL 5.2, 11.1, 13.1, 15.1 and 25.2). Four QTL were detected by both SNP genotyping technologies (QTL 3.2, 5.1, 10.1 and 25.1). The overall better performance of RAD-SNPs than Chip-SNPs in detecting QTL associated with BCWD resistance in the NCCCWA population may be due to sample ascertainment bias effects considering that the 57K SNP Chip was developed using a collection of samples from different rainbow trout populations to maximize SNP polymorphism and discovery (Palti *et al.* 2015a). Therefore, the polymorphic markers in the SNP Chip might be less informative than the SNPs we genotyped with the RAD technology, which were specifically discovered in the sampled families from the NCCCWA population, and are therefore more informative for characterizing genome loci polymorphisms in this dataset.

### Comparing BayesB and wssGBLUP models

Overall, we noticed that the wssGBLUP detected higher number of windows (54) associated with BCWD resistance than the BayesB (20) across the three datasets (TLUM-SNP, NCCCWA-SNP and NCCCWA-RAD) we used here (Table S1). We also noticed that the QTL detection power of BayesB was more negatively impacted by sample size reduction than wssGBLUP: BayesB detected 14, 2 and 4 QTL windows and wssGBLUP detected 31, 9 and 14 QTL windows in TLUM-SNP, NCCCWA-SNP and NCCCWA-RAD datasets, respectively. Performing GS with the Chip genotyped SNPs, we have shown that BayesB predicts GEBVs with higher accuracy than wssGBLUP when using a training sample size of *n* = 1473 (Vallejo *et al.* 2017); however, wssGBLUP outperforms BayesB when using a smaller training sample size of *n* = 583 (Vallejo *et al.* 2016). So, in agreement with these previous GS results, it seems also that the QTL detection power of GWAS with BayesB is more sensitive to sample size reduction than wssGBLUP. We think that the difference in power robustness to sample size reduction of these GWAS models is due to their algorithmic differences. The BayesB model does not explicitly use the available pedigree information and includes in the analysis only animals that had both genotype and phenotype records and as a consequence is less power robust to sample size reduction and needs a minimum sample size for optimal performance, i.e. *n* ≥ 1000 animals. In contrast, the power of GWAS analysis with wssGBLUP is more robust to sample size reduction because it uses the available pedigree information and includes in the analysis animals that have both phenotype and genotype records, and also animals that have only phenotype records (those with missing genotype data).

In the Chip-SNP genotyped TLUM sample, four QTL (3.2, 8.1, 13.2 and 25.1) were detected with both GS models, and four QTL (3.1, 3.3, 5.1 and 13.1) were detected only with wssGBLUP (Table S1). Similarly, in the Chip-SNP genotyped NCCCWA sample, only the QTL 3.2 was detected with both GS models, and most of the QTL (5.1, 10.1, 22.1 and 25.1) were detected only with wssGBLUP. In contrast, in the RAD-SNP genotyped NCCWA sample, three QTL (3.2, 15.1 and 25.1) were detected with both GS models, the QTL 5.2 was detected only with BayesB, and five QTL (5.1, 10.1, 11.1, 13.1 and 25.2) were detected only with wssGBLUP. Thus, these results highlight the importance of using at least two different GWAS algorithms to efficiently uncover the underlying genetic basis of resistance against BCWD in the studied populations.

### Comparing QTL detected in TLUM and NCCCWA populations

In spite of the smaller sample size of the NCCCWA population in comparison to the TLUM population, we detected 10 QTL in the NCCCWA population and only eight QTL in the TLUM population (Table S1). We hypothesize that because the TLUM sample size was much larger than the NCCCWA sample size, it is likely that the type I error rate was smaller in the TLUM sample than in the NCCCWA sample. Therefore, we predict that most of the QTL detected in the TLUM population are real, but some of the QTL detected in the NCCCWA population are false positives. Specifically, the NCCCWA population had two unique QTL (5.2 and 15.1) that were also not reported in past studies. Thus, those two NCCCWA-specific QTL may very be false positives.

### Comparing our GWAS results with previous studies

Eleven of the 14 QTL we detected in this study were also reported in previous studies in the NCCCWA germplasm and in other populations (Table S7) (Johnson *et al.* 2008; Wiens *et al.* 2013; Campbell *et al.* 2014; Quillet *et al.* 2014; Vallejo *et al.* 2014a; Vallejo *et al.* 2014b; Liu *et al.* 2015b; Palti *et al.* 2015b). Campbell et al. (Campbell *et al.* 2014) detected RAD SNPs associated with BCWD resistance in another commercial rainbow trout population that were about 1Mb from our QTL Omy8.1 (Table S1 and Table S7); they also reported QTL that overlap or are close to our detected QTL 10.1 and 25.2. Kutyrev et al. (Kutyrev *et al.* 2016) measured the expression of immune relevant genes on spleen tissue sampled from BCWD resistant (ARS-Fp-R) and susceptible (ARS-Fp-S) genetic lines after laboratory disease pathogen challenge and detected differential expression between the tested genetic lines for genes *il1r-like-1* and *tnfrsf1a-like-a*. Interestingly, two SNPs for the gene *il1r-like-1* (Affx-88933101 and Affx-88915186) were about 240Kb from our QTL 3.2 which explained up to 5.6% of the genetic variance in the NCCCWA population.

Previous studies also detected QTL for BCWD resistance on chromosomes Omy1 (Vallejo *et al.* 2014a; Palti *et al.* 2015b), 2 (Vallejo *et al.* 2014a; Liu *et al.* 2015b), 7 (Quillet *et al.* 2014; Vallejo *et al.* 2014a; Palti *et al.* 2015b), 12 (Vallejo *et al.* 2014a; Liu *et al.* 2015b), 17 (Johnson *et al.* 2008; Campbell *et al.* 2014; Quillet *et al.* 2014), 26 and 28 (Liu *et al.* 2015b) which were not detected in this study. These conflicting results in QTL mapping can be expected due to several reasons including: (1) QTL effects can be population and/or family specific with unique extent/phase of linkage between QTL and marker alleles; and (2) they can also represent false positive results due to limitations and weaknesses of experimental-design and power of analysis as we describe here.

### Conclusion

This GWAS is the most comprehensive genome-wide scan for QTL associated with BCWD resistance performed to date in two commercially-relevant rainbow trout breeding populations, using two whole-genome SNP genotyping platforms and two multiple-regression GWAS models. We identified a total of 14 moderate-large effect QTL associated with resistance to BCWD resistance, and four of those QTL were segregating in the two populations. These GWAS results confirmed that the genetic architecture of BCWD resistance is controlled by the oligogenic inheritance of few moderate-large effect genes and many small effect resistance loci. Overall, the wssGBLUP detected higher number of QTL than the BayesB and both GWAS models did not detect the same QTL which highlights the utility of using two different GWAS algorithms to effectively discover QTL. The RAD genotyping platform detected higher number of QTL than the Chip technology and also both genotyping platforms did not detect the same QTL in the NCCCWA population. These GWAS results will advance the biological and functional analysis of positional candidate genes using the annotation of the new rainbow trout reference genome (GenBank assembly Accession GCA_002163495). They are also likely to be used in the implementation of more efficient selective breeding strategies which will utilize the QTL-flanking SNPs in genome-enabled selection for BCWD resistance in rainbow trout aquaculture.

## ACKNOWLEDGEMENTS

This project was supported by funds from the USDA Agricultural Research Service in house project numbers 8082-32000-006 and 8082-31000-012. We would like to acknowledge the following people for providing technical assistance including Roseanna Long, Kristy Shewbridge, and Cassandra Parker for sample preparation and genotyping; Caird Rexroad, Josh Kretzer, Kyle Jenkins, Travis Moreland, Clayton Birkett, Jen and Ryan Lipscomb and Bryce Williams for fish rearing, phenotyping and sampling. The authors are very grateful to Ignacy Misztal, Shogo Tsuruta, Rafael Silva and Daniela Lourenco for insightful discussions on performing single-step GBLUP analysis with software BLUPF90. We also acknowledge Dorian Garrick for helping to perform GWAS analysis using Bayesian methods with software GENSEL. Mention of trade names or commercial products in this publication is solely for the purpose of providing specific information and does not imply recommendation or endorsement by the U.S. Department of Agriculture. USDA is an equal opportunity provider and employer.

## SUPPLEMETAL MATERIAL

### Supplemental Figures

**Figure S1.**
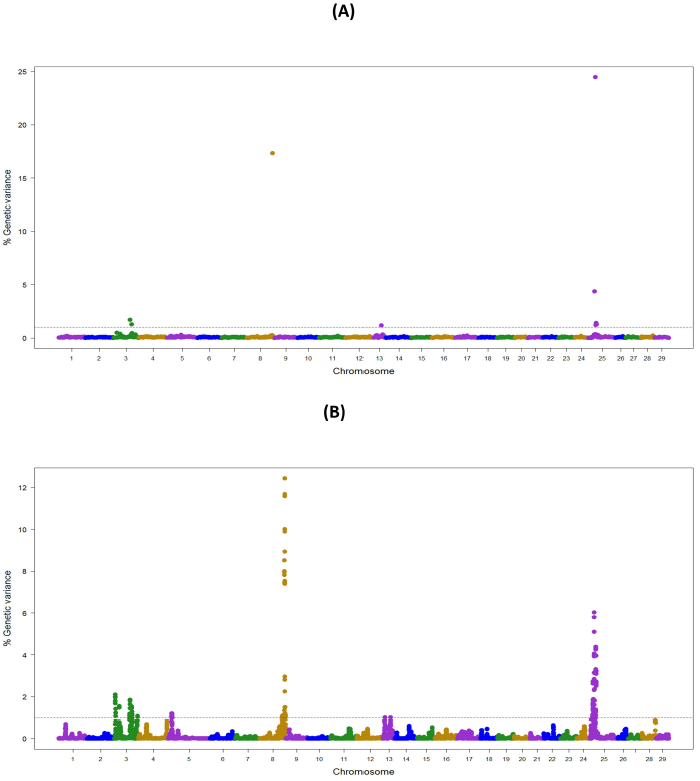
Manhattan plot showing the association between SNP genomic windows and BCWD resistance in TLUM sample genotyped with 57K Chip-SNP: (A) GWAS for DAYS performed with BayesB using 1Mb exclusive windows. (B) GWAS for DAYS performed with wssGBLUP using 1Mb sliding windows. (PPTX file)

**Figure S2.**
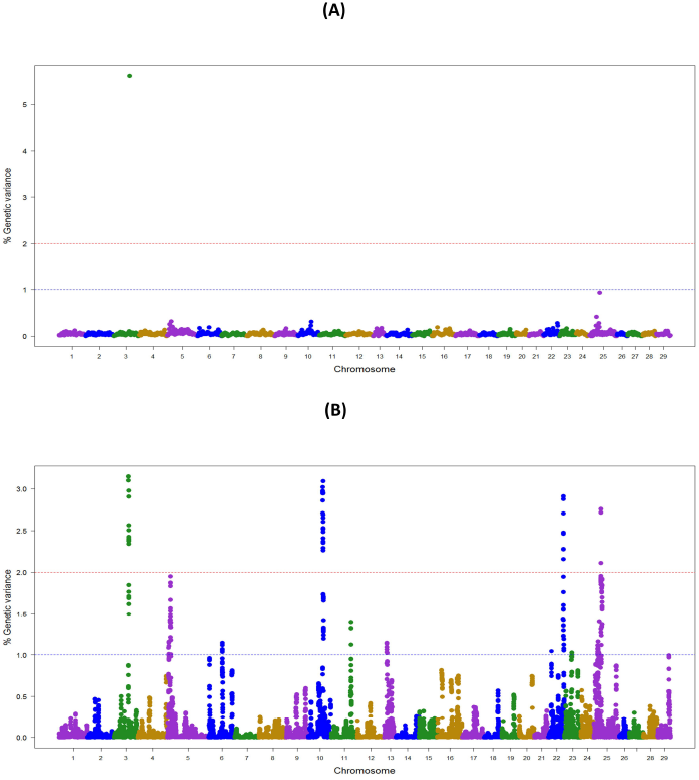
Manhattan plot showing the association between SNP genomic windows and BCWD resistance in NCCCWA sample genotyped with 57K Chip-SNP: (A) GWAS for DAYS performed with BayesB using 1Mb exclusive windows. (B) GWAS for DAYS performed with wssGBLUP using 1Mb sliding windows. (PPTX file)

**Figure S3.**
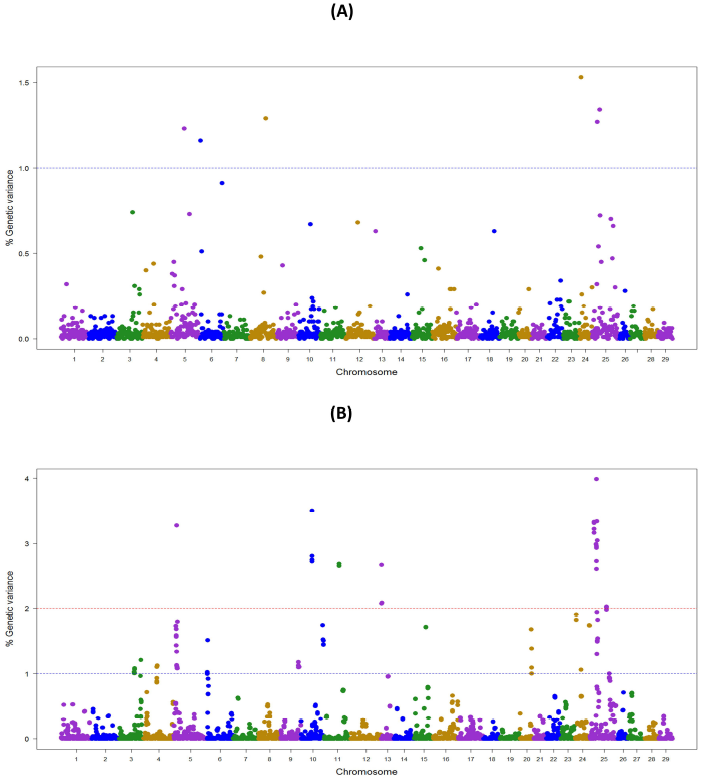
Manhattan plot showing the association between SNP genomic windows and BCWD resistance in NCCCWA sample genotyped with the RAD-SNPs: (A) GWAS for DAYS performed with BayesB using 1Mb exclusive windows. (B) GWAS for DAYS performed with wssGBLUP using 1Mb sliding windows. (PPTX file)

### Supplemental Tables

**Table S1**

Summary of QTL associated1 with BCWD resistance detected using two GWAS methods and two SNP genotyping platforms in NCCCWA and TLUM populations. (XLSX file)

**Table S2**

Summary of QTL associated with BCWD survival DAYS in the Troutlodge US May (TLUM) population. (DOCX file)

**Table S3**

Summary of QTL associated with BCWD survival DAYS in NCCCWA population detected using the 57K Chip-SNP. (DOCX file)

**Table S4**

Summary of QTL associated with BCWD survival DAYS in NCCCWA population detected using RAD-SNPs genotyping. (DOCX file)

**Table S5**

QTL for BCWD resistance that are shared or segregating in both NCCCWA and TLUM populations. (XLSX file)

**Table S6**

QTL for BCWD resistance that are private to either NCCCWA or TLUM population. (XLSX file)

**Table S7**

Summary of QTL for BCWD resistance reported in previous studies in rainbow trout populations. (XLSX file)

